# Alcohol intake triggers aberrant synaptic pruning leading to synapse loss and anxiety-like behavior

**DOI:** 10.1101/870279

**Authors:** Renato Socodato, Joana F. Henriques, Camila C. Portugal, Tiago O. Almeida, Joana Tedim-Moreira, Renata L. Alves, Teresa Canedo, Cátia Silva, Ana Magalhães, Teresa Summavielle, João B. Relvas

**Affiliations:** Instituto de Investigação e Inovação em Saúde and Instituto de Biologia Molecular e Celular, Universidade do Porto. Porto, Portugal

## Abstract

Alcohol use adversely impacts the life of millions of people worldwide. Deficits in synaptic transmission and in microglial function are common findings in human alcohol users and in animal models of alcohol intoxication. Here, we show that alcohol intake over ten consecutive days resulted in substantial loss of excitatory synapse in the prefrontal cortex, a consequence of aberrant synaptic pruning, which led to increased anxiety-like behavior. Mechanistically, these effects of alcohol intake were mediated by a detrimental increase of microglia engulfment capacity via Src-dependent activation of NFkB and consequent TNF production. Accordingly, pharmacological blockade of Src activation or TNF production by microglia, genetic ablation of TNF, or diphtheria toxin-mediated conditional ablation of microglia attenuated aberrant synaptic pruning preventing excitatory synapse loss and anxiety-like behavior. Overall, our data suggest that aberrant pruning of excitatory synapses by microglia might disrupt synaptic transmission during alcohol use.

## Introduction

Alcohol use is associated with pathophysiological changes in the brain and in peripheral organs often resulting in life-threatening conditions. Central nervous system (CNS) exposure to alcohol often leads to behavioral deficits (including anxiety, cognitive decline and motor dysfunction) and impairment of synaptic function, a major hallmark of alcohol use, likely underlies such behavioral deficits. Indeed, alcohol detrimentally impacts the pre and post-synaptic compartments and the secretion/recycling of neurotransmitters, ultimately leading to the disruption of excitatory and inhibitory neurotransmission [1, 2]. This detrimental effect of alcohol on synapses can be contributed by a well-established action of alcohol on neurons and potentially through an underappreciated action of alcohol on glial cells [3].

Microglia, the major innate immune cell population in the brain [4], maintain nervous tissue homeostasis, surveilling the CNS parenchyma by continuously extending and retracting their cellular processes, monitoring for tissue damage or infections and checking the functional status of synapses [5]. Following CNS tissue damage or infection, microglia become activated, changing their morphology (into a more ameboid shape), phagocytic capacity and transcriptional profile in order to restore tissue homeostasis [5]. In many neuropsychiatry disorders, however, microglia immune function becomes dysregulated, often leading to overproduction of inflammatory mediators and exacerbated phagocytic activity, which can be detrimental to synapses and behavior [6, 7].

In human alcohol abusers [8] and in animal models of alcohol intoxication [9] alteration of microglial function is associated with neuroimmune activation [10, 11] and might be directly involved in some of the neurotoxic and adverse behavioral effects of alcohol intake [12]. Indeed, it is thought that innate immune system activation contributes to neuroadaptations in different brain regions that are linked with escalating alcohol consumption, tolerance, dependence and relapsing [13, 14]. Consistent with a neuroimmune hypothesis for alcohol addiction [11], knocking out different genes involved in immune responses, which in the brain are exclusively expressed or highly enriched in microglia, decreases voluntary alcohol intake in mice.

In the present work, we demonstrate that ten consecutive days of alcohol intake upregulates TNF signaling in prefrontal cortex microglia enhancing their engulfment capacity, which leads to aberrant synaptic pruning culminating in synapse loss. Overall, our data suggest that aberrant synaptic pruning by microglia might play an important role in the synaptic transmission deficits elicited by alcohol use.

## Results

### Alcohol intake produces a microglia-driven neuroimmune response in the prefrontal cortex

We studied the effect of alcohol intake on microglia by administrating ethanol (EtOH) or water (H_2_O) for 10 consecutive days to adult Cx3cr1^EYFP-CreER/+^ mice (**Fig. 1A**). Immunohistochemistry on prefrontal cortex tissue sections from Cx3cr1^EYFP-CreER/+^ mice revealed that this EtOH exposure protocol induced a significant expansion of prefrontal cortex microglia (YFP^+^ cells, **Fig. 1B**). Using flow cytometry we confirmed this increased microglia number in Cx3cr1^EYFP-CreER/+^ mice exposed to EtOH (**Fig. 1C**). EtOH also increased the expression of the reactivity markers CD45, integrin alpha M (CD11b) and Iba1 in prefrontal cortex microglia (**Fig. 1C and D**). Morphometry of prefrontal cortex microglia (labeled with Iba1) using Sholl analysis revealed that EtOH induced microglia hyperamification (increased branching (intersections) and process number without affecting process length; **Fig. 1E**). In addition, EtOH significantly altered the expression levels of the microglia-enriched mRNA transcripts P2ry12, Pu.1, Gpr34, Csfr1, C1qC, Tgfβr1, Mertk, Tlr4, Trem2 and C1qA, in the neocortex of Cx3cr1^EYFP-CreER/+^ mice (**Fig. 1F**). Enrichment analysis based on gene ontology using those altered transcripts revealed that several immune-associated pathways were significantly modified by EtOH intake, including pathways controlling microglial activity, innate immune response, and TNF/NFkB signaling (**Fig. 1G**).

**Figure 1.**
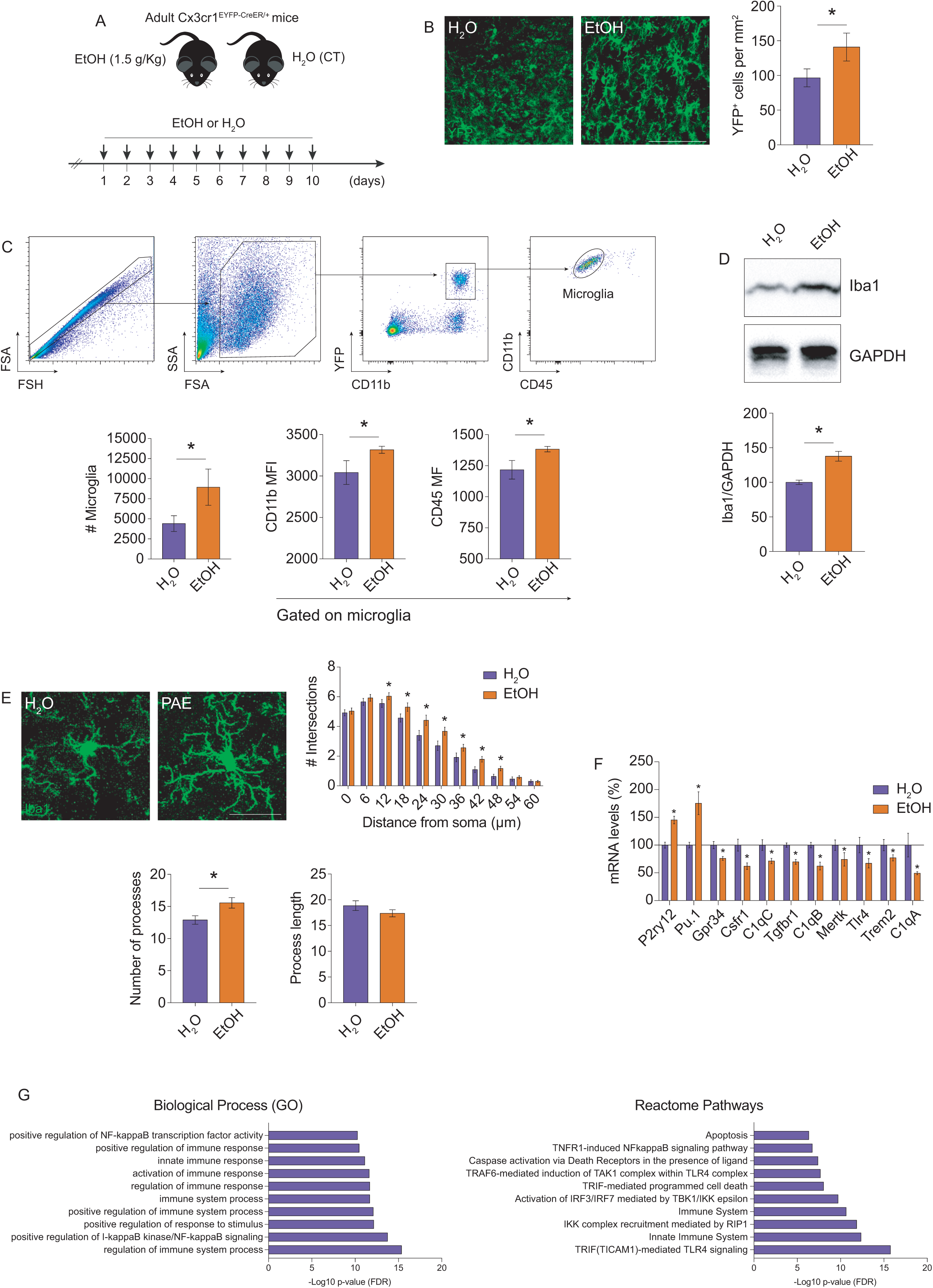
Alcohol intake elicits microglia activation in the prefrontal cortex. **A**, regimen of alcohol (EtOH) or water (H_2_O) exposure (oral gavage for 10 consecutive days) to adult Cx3cr1^EYFP-CreER/+^ mice. **B,** histological confocal analysis for YFP on tissue sections from prefrontal cortices of Cx3cr1^EYFP-CreER/+^ mice exposed to EtOH or H_2_O (n=6 animals per group). Scale bar: 100 μm. **C**, Flow cytometry analysis of microglial numbers in neocortices of Cx3cr1^EYFP-CreER/+^ mice exposed to EtOH or H_2_O (n=8-9 animals). Microglia were identified as being CD11b^+^ and Cd45^low/mid^ gated from the double-positive CD11b^+^ YFP^+^population. **D**, Western blot for Iba1 on lysates from prefrontal cortices of Cx3cr1^EYFP-CreER/+^ mice exposed to EtOH or H_2_O (n=6 animals per group). GAPDH (loading control). **E,** Sholl analysis retrieved from histological confocal images of Iba1 on tissue sections from prefrontal cortices of Cx3cr1^EYFP-CreER/+^ mice exposed to EtOH or H_2_O (n=50 cells from 5 animals per group for each time point). Scale bar: 20 μm. **F,** qRT-PCR from neocortices of Cx3cr1^EYFP-CreER/+^ mice exposed to EtOH or H_2_O (n=5 animals per group). The clustered histogram shows the Z scores of mRNA transcripts normalized to the H_2_O values. **G**, Pathway enrichment analysis inputting the transcripts shown in F. Graphs show the top 10 most overrepresented ontology terms in the GO biological process and in the Reactome pathways. Data on graphs are mean ± SEM. *P<0.05 (Mann-Whitney test in B-D, F; Two-way ANOVA in E).

The effect of EtOH in affecting microglia was also confirmed in wild-type mice, displaying similar results compared with Cx3cr1^EYFP-CreER/+^ mice that carry only one functional Cx3cr1 allele (**Suppl. Fig. 1A and B**). We then evaluated the effect of alcohol on other immune cell populations. Compared with water-treated controls, EtOH did not alter the number of macrophages (EYFP^+^CD11b^+^CD45^hi^ cells), B lymphocytes (CD19^+^CD45^+^CD11b^-^EYFP^-^ cells) or T lymphocytes (CD3^+^CD45^+^CD11b^-^EYFP^-^ cells) in the neocortex of Cx3cr1^EYFP-CreER/+^ mice (**Suppl. Fig. 1C-E**). Astrocytes also perform immune cell functions within the brain parenchyma and are responsive to alcohol. However, we found no significant difference in the number of prefrontal cortex astrocytes (GFAP^+^ cells) or in the expression of GFAP in EtOH-treated mice (**Suppl. Fig. 1F and G**). Collectively, these data indicate that microglia are the major immune cell population affected by EtOH in the prefrontal cortex.

Because alcohol triggered microgliosis, we evaluated the transcript abundance of genes involved in classical immune responses and antioxidant defenses in mice exposed to EtOH. We found that EtOH upregulated the transcript levels of Tlr2, CD14, Icam1, Slc23a2, Tspo, TNF, Rip1, Traf2 and downregulated the transcript levels of IL-6, Ccl5, Tlr7, Mhc-II, Gclc, Traf1, Ccl2, Hmox1, Cox2, Irak3, Tnxnrd1 and Gsr in the neocortex of Cx3cr1^EYFP-CreER/+^ mice (**Fig. 2A**). Moreover, other immune-related genes such as IL-18, IL1b, Inf-b, CD163, Socs3, Nrlp3, Nos2, Cx3cl1, Cxcl10, Arg1, Mrp8, Mrp14, Cxcl1 and Spp1 were not significantly altered by EtOH (**Fig. 2B**). Pathway analysis of the differentially expressed genes following EtOH exposure revealed a substantial number of significantly enriched pathways that were both directly and indirectly modulated by TNF (**Fig. 2C**). The majority of those enriched pathways were related with immune responses but also included cell migration, cell proliferation, cell adhesion, intracellular signal transduction and cytokine biosynthesis/secretion (**Fig. 2C**). Network analysis of protein-protein interactions revealed the prevalence of ten major protein clusters to be modulated by EtOH intake (**Fig. 2D**). The TNF cluster was more intricately associated with Tlr2 and Irak3 clusters, suggesting that the alteration of neuroimmune pathways following EtOH exposure might be co-regulated via TNF/Tlr2/Irak3 (**Fig. 2D**). In addition, several interacting proteins within the TNF cluster (highlighted in blue; **Fig. 2D**), are known modulators/integrators of brain immune responses and microglial activation, thereby indicating that the neuroimmune response triggered by EtOH intake might be strongly associated with TNF.

**Figure 2.**
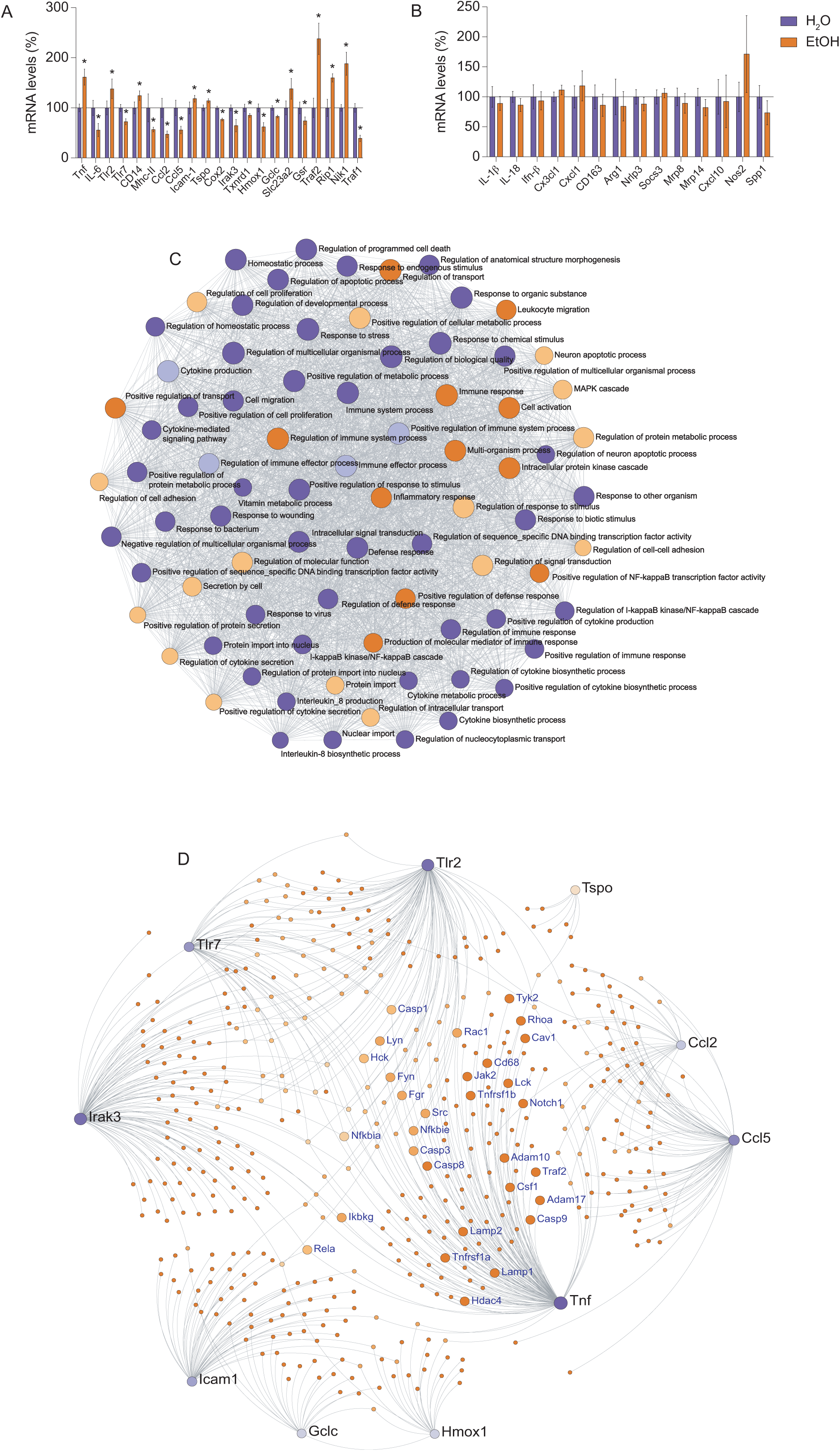
Alcohol intake triggers a TNF-associated neuroimmune response. **A and B,** qRT-PCR from neocortices of Cx3cr1^EYFP-CreER/+^ mice exposed to EtOH or H_2_O (n=5 animals per group). The clustered histogram shows the Z scores of mRNA transcripts normalized to the H_2_O values. Transcripts differentially altered by EtOH are shown in A. Transcripts shown in B were not significantly altered by EtOH. **C**, Pathway enrichment analysis inputting the transcripts shown in A. The topographic pathway map shows overrepresented terms and the degree of interaction between each node. All nodes displayed in the map are modulated directly or indirectly through TNF signaling. **D**, String-based protein-protein interaction (PPI) networks constructed inputting the transcripts shown in A. The PPI network shows the 10 protein clusters to be significantly modulated by EtOH. Substantial overlap occurs between TNF, Irak3 and Tlr2 clusters. Proteins highlighted within the TNF cluster (blue) are known to directly modulate inflammatory responses and immune cell function. Data on graphs are mean ± SEM. *P<0.05 (Mann-Whitney test in A and B).

### Preventing TNF production suppresses microglia activation following alcohol intake

To test whether microglia were the major producers of TNF during EtOH exposure we genetically ablated microglia using the microglial iDTR/Cre-lox system [15]. We first validated the efficiency of this microglia ablation system in the context of our experimental requirements by giving tamoxifen to Cx3cr1^EYFP-CreER/+^:R26^iDTR/+^ and Cx3cr1^EYFP-CreER/+^ mice at P28 and P30 and diphtheria toxin (DT) 2 months later (**Fig. 3A**). Indeed, both immunohistochemistry on prefrontal cortex tissue sections and flow cytometry showed, as expected, that microglia were efficiently eliminated in Cx3cr1^EYFP-CreER/+^:R26^iDTR/+^ mice (**Fig. 3B and C, respectively**).

**Figure 3.**
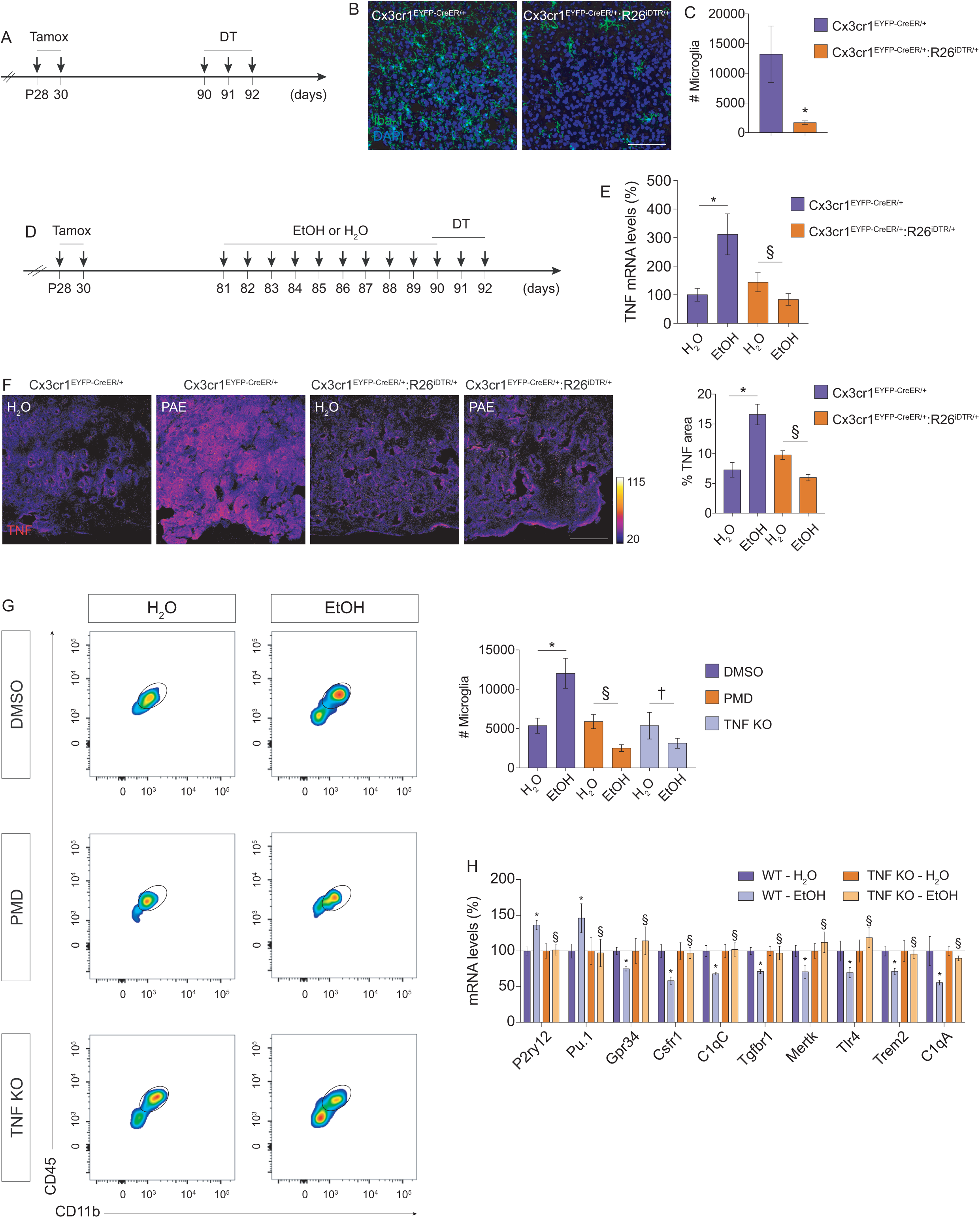
TNF production elicited by alcohol intake drives microglia activation in the prefrontal cortex. **A**, scheme for tamoxifen-inducible microglial ablation in the brain. **B,** histological confocal analysis for Iba1 on tissue sections from prefrontal cortices of Cx3cr1^EYFP-CreER/+^ and Cx3cr1^EYFP-CreER/+^:R26^iDTR/+^ after exposure to diphtheria toxin (DT; n=4 animals per genotype). Scale bar: 50 μm. **C**, flow cytometry analysis of microglial numbers in neocortices of Cx3cr1^EYFP-CreER/+^ and Cx3cr1^EYFP-CreER/+^:R26^iDTR/+^ after exposure to DT (n=6 animals per genotype). **D**, scheme for microglial ablation during EtOH exposure. **E,** qRT-PCR from neocortices of Cx3cr1^EYFP-CreER/+^ and Cx3cr1^EYFP-CreER/+^:R26^iDTR/+^ after exposure to EtOH or H_2_O and treatment with DT (n=5 animals per genotype/group). Histogram shows the Z scores of TNF transcripts normalized to the Cx3cr1^EYFP-CreER/+^ - H_2_O values. **F,** histological confocal analysis for TNF on tissue sections from prefrontal cortices of Cx3cr1^EYFP-CreER/+^ and Cx3cr1^EYFP-CreER/+^:R26^iDTR/+^ after exposure to EtOH or H_2_O and treatment with DT (n=5 animals per genotype). Scale bar: 100 μm. **G**, flow cytometry analysis of microglial numbers in neocortices of wild-type (WT) or TNF knockout (KO) mice exposed to EtOH or H_2_O (n=8-10 animals per group). WT animals were also treated with DMSO or with pomalidomide (PMD). **H,** qRT-PCR from neocortices of WT or TNF KO mice exposed to EtOH or H_2_O (n=5 per group). Histogram shows the Z scores of mRNA transcripts normalized to the WT - H_2_O values. Data on graphs are mean ± SEM. *P<0.05 (Mann-Whitney test in C; Two-way ANOVA E-H). ^§^not significantly different, ^†^not significantly different (Two-way ANOVA).

Next, we evaluated the production of TNF in the prefrontal cortex of Cx3cr1^EYFP-CreER/+^ and microglia-depleted (Cx3cr1^EYFP-CreER/+^:R26^iDTR/+^) mice following EtOH exposure (**Fig. 3D**). Whereas EtOH significantly increased the mRNA levels and the protein amounts of TNF in Cx3cr1^EYFP-CreER/+^ mice (**Fig. 3E and F, respectively**), it did not increase the amounts of TNF in microglia-depleted mice (**Fig. 3E and F**), suggesting that microglia are the major producers of TNF in the prefrontal cortex during EtOH exposure.

We next used TNF-deficient (TNF KO) mice or the immunomodulatory and brain-penetrant TNF blocker pomalidomide (PMD) [16–18] to test whether TNF production could lead to microgliosis. Using flow cytometry, we found that the EtOH-induced microgliosis (**Fig. 3G**) was completely prevented in TNF KO mice (**Fig. 3G**) or in wild-type mice treated with PMD (**Fig. 3G**). The EtOH-induced alteration of the microglia enriched genes P2ry12, Pu.1, Gpr34, Csfr1, C1qC, Tgfβr1, Mertk, Tlr4, Trem2 and C1qA was also abrogated in TNF KO mice (**Fig. 3H**), indicating that TNF produced by microglia during EtOH exposure is the trigger for microgliosis.

### Alcohol intake modulates Src tyrosine kinase/NFκB pathway to increase TNF production and trigger microglia activation

Because the tyrosine kinase Src controls microglial activation via increased production and secretion of TNF [19–21], we investigated whether EtOH could trigger the production of TNF in microglia via Src. We found that the amounts of active Src (phospho-Src Tyr^146^) were significantly increased in the prefrontal cortex of Cx3cr1^EYFP-CreER/+^ mice exposed to EtOH (**Fig. 4A**). However, the amounts of active Src (phospho-Src Tyr^146^) were not significantly increased in the prefrontal cortex of microglia-depleted mice exposed to EtOH (**Fig. 4A**). In line with this, the effect of alcohol in increasing microglial Src activation was thoroughly reproduced in a cell autonomous manner in primary cortical microglia (analyzed by immunocytochemistry with an antibody against phospho-Src Tyr^146^ (**Suppl. Fig. 2A**)) and in the CHME3 microglial cell line (analyzed by live cell imaging using a FRET-based Src biosensor (**Suppl. Fig. 2B**)).

**Figure 4.**
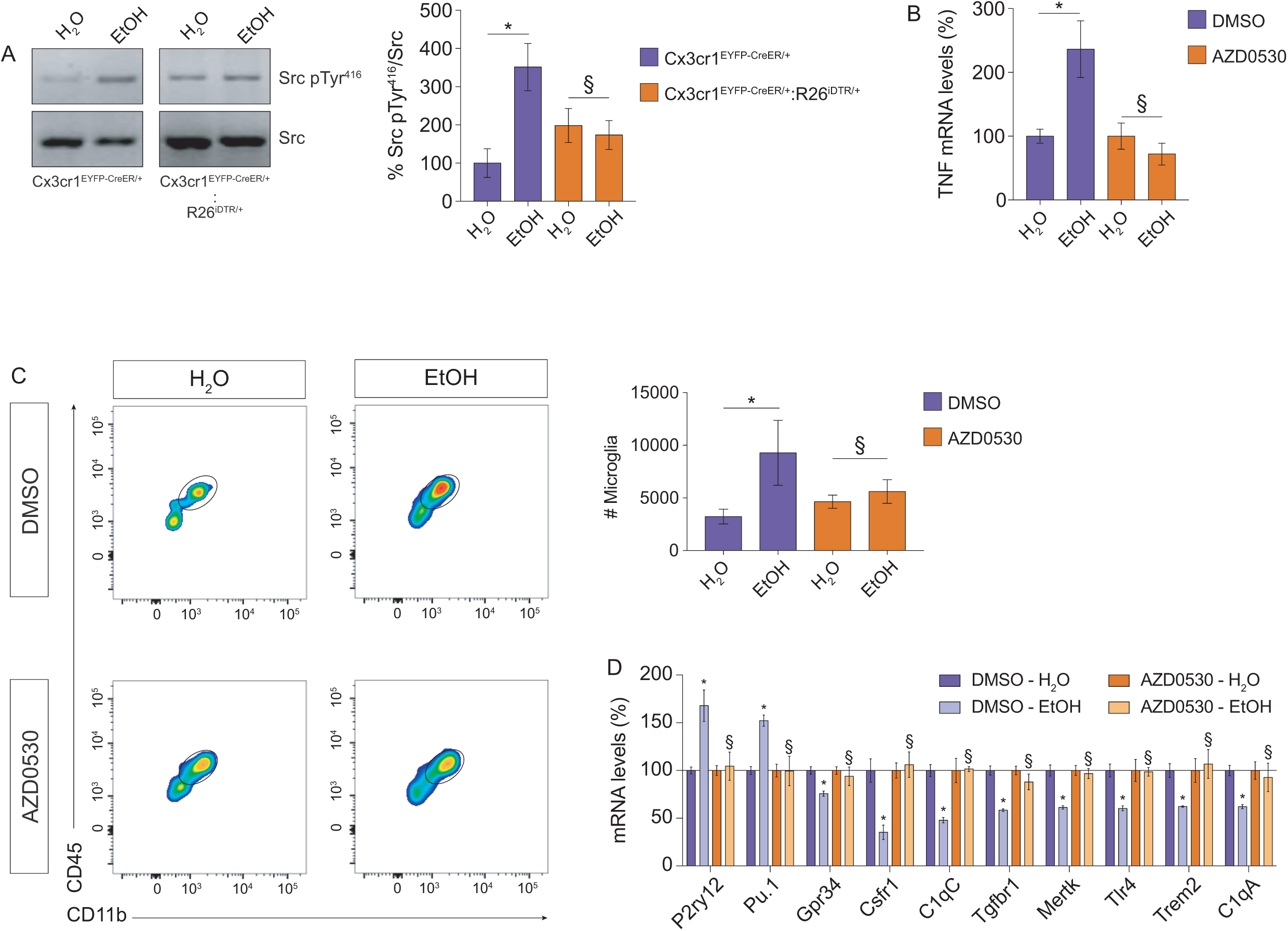
Src activation elicited by alcohol intake leads to TNF production driving prefrontal cortex microglia activation. **A,** Western blot analysis for Src phospho-Tyr^416^ (active form) on lysates from prefrontal cortices of Cx3cr1^EYFP-CreER/+^ and Cx3cr1^EYFP-CreER/+^:R26^iDTR/+^ after exposure to EtOH or H_2_O and treatment with DT (n=5 animals per genotype). Src (loading control). **B**, qRT-PCR from neocortices of Cx3cr1^EYFP-CreER/+^ mice exposed to EtOH or H_2_O (n=5 animals per group). Mice were treated with either DMSO or the brain-penetrant Src inhibitor AZD0530. **C**, flow cytometry analysis of microglial numbers in neocortices of Cx3cr1^EYFP-CreER/+^ mice exposed to EtOH or H_2_O (n=8 animals per group). Mice were treated with either DMSO or AZD0530. **D**, qRT-PCR from neocortices of Cx3cr1^EYFP-CreER/+^ mice exposed to EtOH or H_2_O (n=5 animals per group). Mice were also treated with DMSO or with AZD0530. The clustered histogram shows the Z scores of mRNA transcripts normalized to the DMSO - H_2_O values. Data on graphs are mean ± SEM. *P<0.05, ^§^not significantly different (Two-way ANOVA)

To confirm the relationship between Src activity and TNF expression we used the clinically relevant Src blocker AZD0530 [22, 23] during EtOH exposure. We found that inhibiting Src with AZD0530 (**validation in Suppl. Fig. 2C**) abrogated the increase in TNF elicited by EtOH (**Fig. 4B**). To further confirm that alcohol modulates microglial Src to increase the production of TNF, we inhibited Src, using the Src blocker SKI-1, in primary cortical microglia and found that SKI-1 prevented the alcohol-mediated increase of TNF production (**Suppl. Fig. 2D**).

To gain further mechanistic insight into how Src drives TNF production in microglia during alcohol exposure we focused on the role of NFκB, a major regulator of inflammatory signaling in immune cells [24, 25]. We measured the nuclear accumulation of the p65 subunit of NF-kB as a functional indicator of NF-kB activation in CHME3 microglia using a green fluorescent protein (GFP)–tagged p65 construct. We found that exposure to alcohol increased nuclear GFP content in CHME3 microglia (**Suppl. Fig. 2E**), and that knocking down Src (**validation in Suppl. Fig. 2F**), abrogated the alcohol-induced nuclear translocation of GFP-p65 subunit (**Suppl. Fig. 2E**). Pharmacological blockade of Src using AZD0530 or SKI-1 also prevented the alcohol-induced activation of the NFκB pathway (monitored using the IκBα-miRFP703 reporter [26]; **Suppl. Fig. 2G**) Moreover, direct activation of Src in CHME3 microglia, using rapamycin-based chemogenetics with RapR-Src/FRB [27], increased the nuclear translocation of the GFP-p65 subunit (**Suppl. Fig. 2H**). Paralleling the increase of NFκB activation following Src activation, overexpressing the constitutively active Src mutant Src^Y527F^ led to increased secretion of TNF from microglia (**Suppl. Fig. 2I**). Finally, preventing NFκB activation with the clinically relevant inhibitor sulfasalazine [28, 29] completely blocked the production of TNF elicited by Src^Y527F^ in microglia (**Suppl. Fig. 2J**). These data suggest that alcohol drives TNF production in microglia via Src and downstream NFκB activation.

In line with such Src-dependent TNF production during alcohol exposure, blocking Src with AZD0530 abrogated the EtOH-induced increase of microglial numbers and prevented the EtOH-mediated alteration of the microglia-enriched genes P2ry12, Pu.1, Gpr34, Csfr1, C1qC, Tgfβr1, Mertk, Tlr4, Trem2 and C1qA (**Fig. 4C and D**).

### Blocking microglial TNF signaling suppresses anxiety-like behavior elicited by alcohol intake

The neuroimmune hypothesis of alcohol addiction [11] supports the notion that alterations of neuroimmune signaling might be responsible for some of the behavioral deficits elicited by alcohol abuse. Because EtOH induced the production of TNF by prefrontal cortex microglia, we tested whether EtOH could drive changes in behavior. We found, using the elevated-plus maze (EPM) test, that EtOH increased anxiety-like behavior in adult Cx3cr1^EYFP-CreER/+^ mice, as revealed by the decreased time that EtOH-subjected mice spent on the open arms (**Fig. 5A**). No differences were found in the distance travelled in the EPM between EtOH-treated and water-treated mice (**Fig. 5B**). The anxiogenic effect of EtOH could also be observed in wild-type mice in the EPM test (**Suppl. Fig. 3A and B**). Moreover, the increase of anxiety-like behavior elicited by EtOH exposure was reproduced in the open field test, as mice exposed to EtOH spent less time in the center of the open field arena than water-treated littermates (**Suppl. Fig. 3C**).

**Figure 5.**
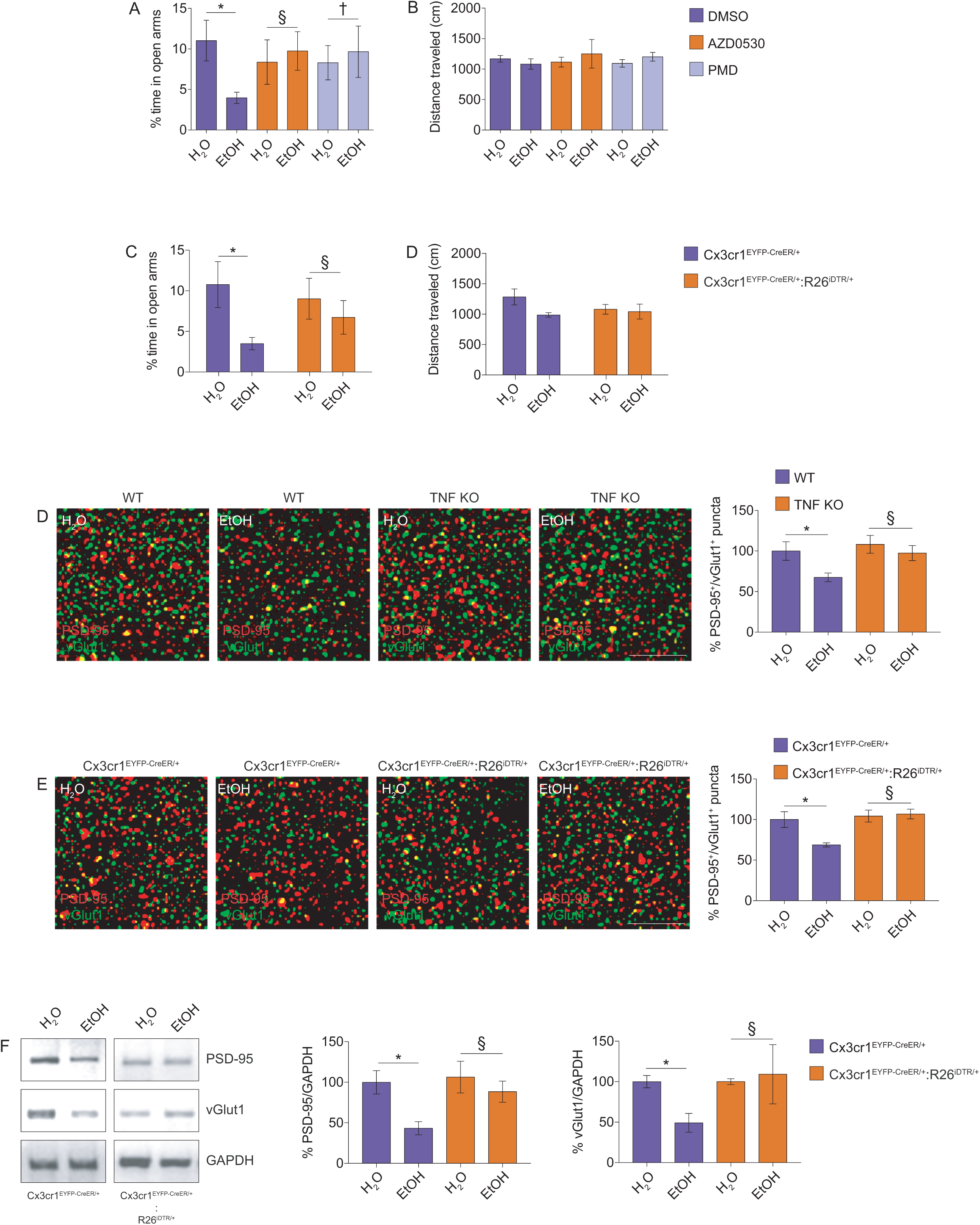
Alcohol intake elicits microglia-dependent synapse loss and anxiety-like behavior. **A-B**, Cx3cr1^EYFP-CreER/+^ mice were injected with DMSO, AZD0530 or PMD and exposed to EtOH or H_2_O and then evaluated in the elevated-plus maze (EPM) test (n=9-13 animals per group). **C-D,** Cx3cr1^EYFP-CreER/+^ and Cx3cr1^EYFP-CreER/+^:R26^iDTR/+^ mice were given tamoxifen, treated with diphtheria toxin, exposed to EtOH or H_2_O and then evaluated in the EPM test (n=6 animals per group). **E,** histological confocal colocalization analysis for PSD-95 (red) and vGlut1 (green) on tissue sections from prefrontal cortices of WT and TNF KO mice after exposure to EtOH or H_2_O (n=6-9 animals per genotype). Scale bar: 5 μm. **F,** histological confocal colocalization analysis for PSD-95 (red) and vGlut1 (green) on tissue sections from prefrontal cortices of in Cx3cr1^EYFP-CreER/+^ and Cx3cr1^EYFP-CreER/+^:R26^iDTR/+^ mice after exposure to EtOH or H_2_O and treatment with DT (n=6-9 animals per genotype). Scale bar: 5 μm. **G,** Western blot analysis for PSD-95 and vGlut1 on lysates from prefrontal cortices of Cx3cr1^EYFP-CreER/+^ and Cx3cr1^EYFP-CreER/+^:R26^iDTR/+^mice after exposure to EtOH or H_2_O (n=5 animals per group). GAPDH (loading control). Data on graphs are mean ± SEM. *P<0.05 (Two-way ANOVA in A-F) ^§^not significantly different, ^†^not significantly different (Two-way ANOVA).

Importantly, EtOH did not produce significant changes in (1) general locomotor activity (**Suppl. Fig. 3D**) or repetitive behavior (**Suppl. Fig. 3E**) when mice were evaluated in the open field test, and in (2) recognition memory when mice were evaluated in the novel object recognition test (**Suppl. Fig. 3F and G**).

We then investigated whether the anxiogenic effect of EtOH intake required the production of TNF by microglia. We found that blocking TNF production indirectly (by inhibiting Src with AZD0530) or directly (by inhibiting TNF expression with PMD) completely suppressed the EtOH-induced increase in anxiety-like behavior (**Fig. 5A-C**). To further confirm the role of microglia in the anxiogenic effect of EtOH intake, we evaluated microglia-depleted mice in the EPM test. Whereas EtOH increased anxiety-like behavior in control (Cx3cr1^EYFP-CreER/+^) mice, this anxiogenic effect was significantly prevented in microglia-depleted (Cx3cr1^EYFP-CreER/+^:R26^iDTR/+^) mice (**Fig. 5C**). No differences were found in the distance travelled in the EPM between the genotypes or treatment groups (**Fig. 5D**).

### TNF signaling increases microglia phagocytic activity licensing microglia to prune prefrontal cortex synapses following alcohol intake

Behavioral alterations, such as increased anxiety, can be caused by perturbations in excitatory/inhibitory balance in prefrontal cortex circuits [30, 31] and microglia activation, elicited by EtOH, could alter excitatory synapses in the prefrontal cortex. Indeed, we found that the number of excitatory synapses, evaluated by double-labeling immunohistochemistry against the post-synaptic protein PSD-95 and the pre-synaptic protein vGlut1, was significantly decreased in prefrontal cortices of mice exposed to EtOH (**Fig. 5E**). The loss of PSD-95^+^/vGlut1^+^ puncta elicited by EtOH intake was completely prevented in TNF KO mice (**Fig. 5E**). Furthermore, the EtOH-induced decrease of PSD-95^+^/vGlut1^+^ puncta was abrogated in microglia-depleted mice (**Fig. 5F**). Paralleling this decrease of excitatory synapse number, EtOH intake also decreased, in a microglia-dependent manner, the amounts of PSD-95 and vGlut1 in the prefrontal cortex (**Fig. 5G**), indicating that TNF production by microglia leads to loss of excitatory synapses in the prefrontal cortex following EtOH intake.

Loss of synapses can be a consequence of increased neuronal cell death. However, we found that the number of neurons (stained with NeuN) in the prefrontal cortex was not significantly different between EtOH-subjected and water-treated mice (**Suppl. Fig. 3H**). Interestingly, the numbers of excitatory synapses in the CA1 region of the dorsal hippocampus of EtOH-subjected mice were not significantly different from those of water-treated littermates (**Suppl. Fig. 4A**), which may help to explain the specificity of the behavioral phenotype elicited by EtOH.

In pathological conditions, such as in Alzheimer’s disease, microglia can phagocytose and prune healthy synapses, a process termed synaptophagy [7]. We hypothesized that the EtOH-mediated loss of excitatory synapses in the prefrontal cortex could be a direct consequence of excessive engulfment of synaptic structures by microglia. To assess the engulfment of synapses by microglia we prepared synaptosomes (to isolate synaptic terminals) from the prefrontal cortex of adult mice and incubated microglial cultures with them. Indeed, immunofluorescence labeling of PSD-95 confirmed that microglia efficiently engulfed synaptosomes prepared from the prefrontal cortex (**Fig. 6A**), further confirming that microglia actively phagocytose synapses in steady state conditions. However, cultured microglia exposed to EtOH engulfed significantly more prefrontal cortex synaptosomes than control microglia (**Fig. 6A**). To further corroborate the excessive EtOH-mediated engulfment of synaptic structures, microglia cultures were incubated with synaptosomes prepared from the prefrontal cortex of Thy1-YFP mice. Measuring microglia engulfment capacity by flow cytometry revealed that microglia exposed to EtOH displayed significantly increased engulfment of YFP^+^ synaptosomes compared with control microglia (**Fig. 6B**). Blocking TNF with the neutralizing antibody adalimumab (HUMIRA) completely prevented the engulfment of YFP^+^ synaptosomes elicited by exposure to EtOH (**Fig. 6B**). To investigate whether microglia exposed to EtOH engulfed more synapses in vivo, the amount of PSD-95^+^ puncta within the cell body of Iba1^+^ microglia was evaluated by immunofluorescence on prefrontal cortex tissue sections. Confocal imaging coupled with 3D cell surface rendering revealed that Iba1^+^ microglia from the prefrontal cortex of mice exposed to EtOH contained significantly more PSD-95^+^ puncta within their cell bodies than prefrontal cortex microglia from mice treated with water (**Fig. 6C**). This microglial engulfment of prefrontal cortex postsynaptic elements required TNF signaling because no significant differences in the engulfment of PSD-95^+^ puncta were observed in TNF KO mice exposed to EtOH (**Fig. 6C**).Conversely, EtOH neither increased PSD-95 engulfment by Iba1^+^ microglia in the CA1 region of dorsal hippocampus (**Suppl. Fig. 4B**) nor triggered the production of TNF in the dorsal hippocampus (**Suppl. Fig. 4C**).

**Figure 6.**
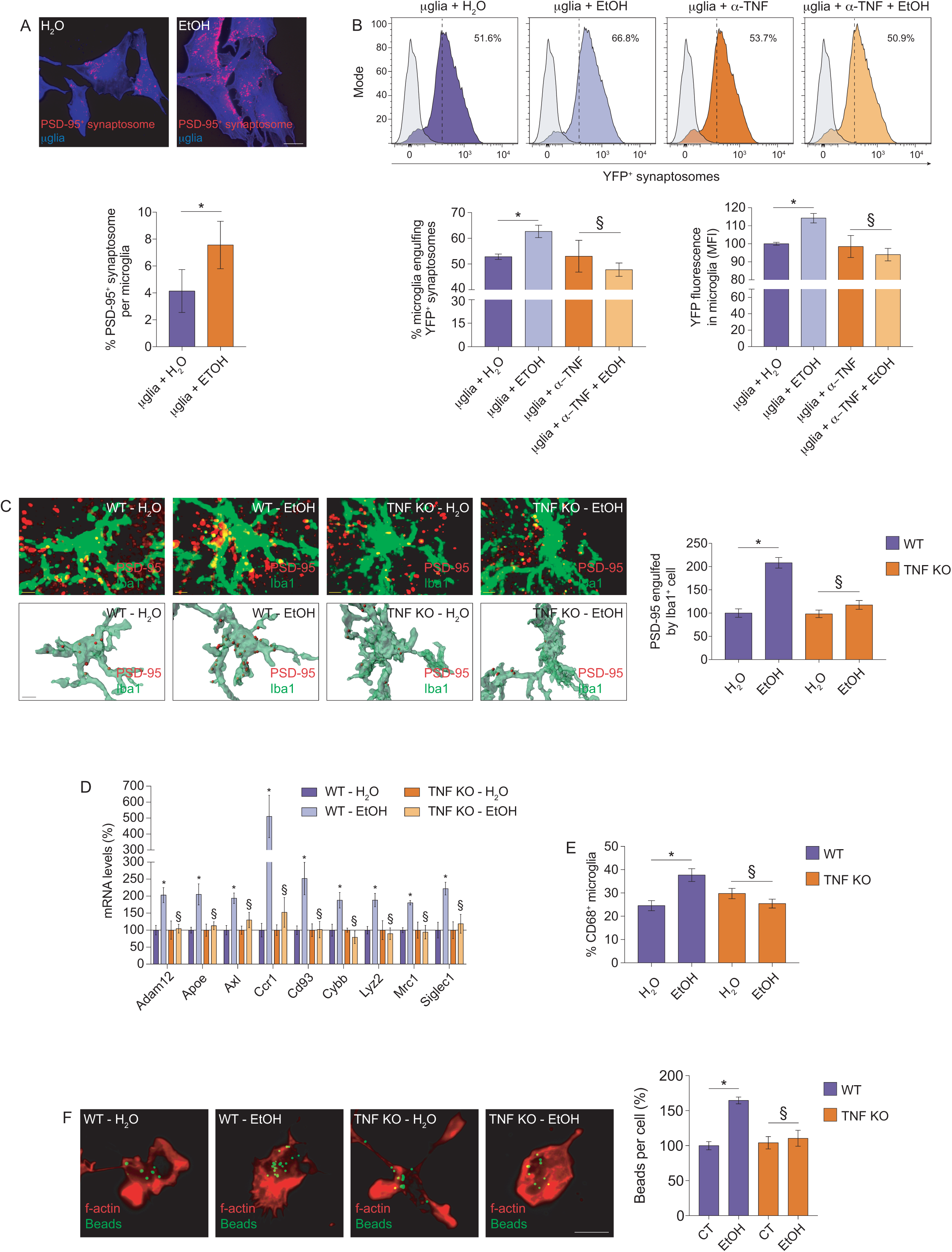
Alcohol intake licenses microglia to prune synapses. **A**, immunofluorescence images of PSD-95-labeled prefrontal cortex synaptosomes (red) inside cultured CHME3 microglia labeled with CellMask dye (blue). Images are representative of 3 different experiments. Scale bar: 20 μm. **B**, representative flow cytometry profiles for YFP labeling in CHME3 microglia incubated for 24 h with synaptosomes isolated from the prefrontal cortex of Thy1-YFP mice (n=6 cultures per group). In some conditions microglial cultures were pretreated for 24 h with EtOH (70 mM) or with the TNF blocking antibody adalimumab (5 μg/ml). Engulfment ability of microglia was calculated by comparing the percentage (left graph) or the MFIs (right graph) of microglia expressing high YFP signal. **C,** representative confocal maximum projection images and relative Imaris 3D surface rendering showing volume reconstruction of PSD-95 engulfed by microglia (Iba1^+^ cell) on tissue sections from prefrontal cortices of WT and TNF KO mice after exposure to EtOH or H_2_O (n=9 animals per group). Scale bar: 3 μm. **D,** qRT-PCR from neocortices of WT and TNF KO mice exposed to EtOH or H_2_O (n=5 animals per group). The clustered histogram shows the Z scores of mRNA transcripts normalized to the DMSO - H_2_O values. **E**, flow cytometry analysis of CD68 expression in microglia (gated as CD45^mid^CD11b^+^ cell population) from prefrontal cortices of Cx3cr1^EYFP-CreER/+^ mice exposed to EtOH or H_2_O (n=5 animals per group). **F,** primary cortical microglial cultures were treated with EtOH (70 mM) or H_2_O (n=4 different and independent cultures), incubated with fluorescent microbeads (green) for 90 min and stained with phalloidin (red). Data on graphs are mean ± SEM. *P<0.05 (unpaired t test in A) *P<0.05 (One-way ANOVA in B) *P<0.05 (Two-way ANOVA in C-F) ^§^not significantly different (One-way ANOVA (B) or Two-way ANOVA (C-F)).

In line with the view that EtOH intake increases microglia engulfment capacity, mice exposed to EtOH displayed a TNF-dependent upregulation of the microglia engulfment module (represented by transcripts coding for Adam12, Apoe, Axl, Ccr1, CD93, Cybb, Lyz2, Mrc1 and Siglec1) [32] (**Fig. 6D**) and also had a TNF-dependent increase of prefrontal cortex microglia expressing the phagocytic marker CD68 (**Fig. 6E**). In this context, TNF signaling enabled microglial phagocytosis because EtOH increased the phagocytic activity of cortical microglia from wild-type mice but not of cortical microglia from TNF KO mice (**Fig. 6F**). These data suggest that EtOH intake results in aberrant pruning of prefrontal cortex excitatory synapses via microglial TNF signaling.

## Discussion

Neuroinflammation is regarded as a main contributing factor in alcohol-induced brain damage [33–36]. Despite inducing neuroimmune activation and excitatory synapse loss, our EtOH exposure protocol simulating binge with moderate alcohol intake produced neither neuronal cell death nor a classical proinflammatory signature, which is different from studies where binge with heavy alcohol intake triggers a proinflammatory signature with concomitant gliosis [37–39]. In fact, our data are more in line with microarray and transcriptomic studies showing that alcohol intake produces neuroimmune activation without triggering robust proinflammatory response [40–44]. Therefore, the moderate dosage of EtOH we used likely explains the neuroimmune activation without robust proinflammatory signature and neuronal cell loss in the prefrontal cortex.

Gene ontology with pathway enrichment indicated that TNF was the most prominent target modulating neuroimmune activation and microglial responses following EtOH intake. In line with this, using the iDTR/Cre-lox system we eliminated microglia and established that they were the main producers of TNF in the prefrontal cortex during EtOH exposure. Then, using pomalidomide, a clinically relevant brain-penetrant [18] immunomodulatory agent that blocks TNF, or TNF-null mice we further confirmed that inhibiting or preventing TNF production, respectively, abrogated microglia activation and associated changes in the expression of RNA transcripts involved in microglial function following EtOH exposure. Thus, we concluded that EtOH initiated a microglial TNF-dependent immune response in the prefrontal cortex.

Src activation following neuroinflammation triggers the production of proinflammatory mediators by microglia, including TNF [21, 22]. EtOH increased Src activation leading to the nuclear translocation of the p65 subunit of the NFκB complex, which culminated in NFκB-dependent TNF production and release by microglia. Paralleling the effect of the blockade of TNF production with pomalidomide, inhibiting Src with AZD0530 (a clinically relevant Src inhibitor) also prevented microgliosis and the increase of anxiety-like behavior elicited EtOH intake, suggesting that targeting the Src/NFκB/TNF pathway likely overcomes the anxiogenic effect of EtOH. However, the precise signaling mechanism by which alcohol exposure elicits microglial Src activation to impact NFκB-dependent transcription warrants further investigation.

The prefrontal cortex contains glutamatergic excitatory neurons synapsing locally or projecting to distant cortical and subcortical nuclei. Failure to set up/sustain proper excitatory connectivity leads to imbalanced neuronal activity across networks [45], which could explain at least some of the behavioral impairments found in neurological disorders [18]. Accordingly, the loss of prefrontal cortex excitatory synapses elicited by EtOH could be sufficient to disrupt the excitatory/inhibitory balance of prefrontal cortex neurons projecting to anxiety-related centers in subcortical regions, implying that EtOH-induced prefrontal cortex synapse loss might have directly impacted the alcohol-related anxiety-like behavior. In this context, our analyses evaluating anxiety-like behavior were carried out 120 min after the last intake of EtOH, a time frame in which intake of 1.5 g/Kg leads to 100-120 mg/dl of EtOH in the blood of adult mice [46, 47]. Therefore, the increased anxiety-like behavior we found in mice exposed to EtOH likely reflects a more persistent anxious state driven by consecutive EtOH intake rather than withdrawal-induced anxiety.

Microglial clearance activity varies significantly across different brain regions [32]. Because of this regional difference in microglia clearance activity it is conceivable that prefrontal cortex microglia were more prone to engulf and prune synapses than their hippocampal counterparts upon exposure to alcohol. Identifying the precise mechanisms underlying such regional differences warrant future in-depth studies focusing on the characterization of microglial heterogeneity.

The phagocytic engulfment of healthy synapses by microglia can result in synapse loss in the brain [7], and in Alzheimer’s disease it can contribute to early disease pathology [48]. The fact that EtOH licensed microglia to prune prefrontal cortex synapses in the absence of significant alterations in the number of neurons in the prefrontal cortex also suggests that aberrant synaptic pruning by microglia might be an early hallmark associated with neurotoxicity during alcohol use. Furthermore, demonstrating that microglia can engulf prefrontal cortex postsynaptic elements provides important new mechanistic evidence into how the brain immune system might contribute to the impairment of synaptic transmission, a major detrimental consequence of alcohol use.

In summary, intake for 10 consecutive days of moderate amounts of alcohol in adult mice elicited a TNF-dependent enhancement of microglia engulfment capacity, which licensed microglia to prune excitatory synapses in the prefrontal cortex. Such microglia-mediated aberrant synaptic elimination in the prefrontal cortex was associated with increased anxiety-like behavior, could be mitigated by pharmacological blockade of TNF signaling and was completely prevented in TNF null mice or in mice in which microglia was conditionally ablated. Overall, our results support the idea that aberrant synaptic pruning by microglia is, at least in part, responsible for alterations in synaptic transmission caused by alcohol use.

## Experimental Procedures

### Animals

All mice experiments were reviewed by i3S animal ethical committee and were approved by Direção-Geral de Alimentação e Veterinária (DGAV). Animals were maintained in standard laboratory conditions with inverted 12h/12h light dark cycle and were allowed free access to food and water. Mice were housed under specific pathogen-free conditions. Experiments were carried out following the 3Rs ethics and mice were kept on a C57BL/6 background. For the present study, only male mice (16-20 week-old) were used as female mice are more prone to develop alcohol-related inflammation and neuronal damage [49, 50], a confounding factor for assessing alterations on microglia and synaptic function. Furthermore, the use of only one gender minimizes intergroup variability and reduces the overall number of animals used.

B6.129P2(Cg)-*Cx3cr1^tm2.1(cre/ERT2)Litt^*/WganJ mice (herein referred asCx3cr1^EYFP-CreER/+^; The Jackson Laboratory Stock No: 021160; RRID:IMSR_JAX:021160) mice were used to study brain microglia in this work. These mice express a Cre-ERT2 fusion protein and an enhanced yellow fluorescent protein (EYFP) from endogenous Cx3cr1 promoter. EYFP fluorescence is observed in more than 95% of Iba1^+^ microglia in the brain [15].

For microglial cell ablation experiments Cx3cr1^EYFP-CreER/+^ mice were intercrossed with R26^iDTR/+^ (*C57BL/6-Gt(ROSA)26Sor^tm1(HBEGF)Awai^/J*; The Jackson Laboratory Stock No: 007900; RRID:IMSR_JAX:007900) mice. Genotypes of interest (Cx3cr1^EYFP-CreER/+^ (control) and Cx3cr1^EYFP-CreER/+^:R26^iDTR/+^ **(experimental**)) were determined by PCR using primers for R26-iDTR insertion including ROSA26-forward: AAA GTC GCT CTG AGT TGT TAT; ROSA26-reverse: GCG AAG AGT TTG TCC TCA ACC; wild-type reverse: GGA GCG GGA GAA ATG GAT ATG and primers for CreER-EYFP insertion including forward: AAG ACT CAC GTG GAC CTG CT; wild-type reverse: AGG ATG TTG ACT TCC GAG TG; mutant reverse: CGG TTA TTC AAC TTG CAC CA.^+^.

Prof. Rui Applelberg (University of Porto) supplied C57BL/6.TNF knockout (referred herein as TNF KO) mice. TNF KO mice were genotyped by PCR using ATC CGC GAC GTG GAA CTG GCA GAA (forward) and CTG CCC GGA CTC CGC AAA GTC TAA (reverse) primer pair. TNF KO mice display a single band of 2 kb in the PCR gel. TNF deficient mice were generated from hemizygote progenitors and wild-type littermates were used as controls.

Tg(Thy1-cre/ERT2,-EYFP)HGfng/PyngJ (also known as SLICK-H line and herein termed Thy1-YFP; The Jackson Laboratory Stock No: 012708; RRID:IMSR_JAX:012708) mice were maintained as before [51]. These mice have constitutive and exclusive YFP labeling of neurons driven by the endogenous Thy1 promoter [52]. In this work, Thy1-YFP mice were used for studying the engulfment of synapses by microglia in vitro.

### Drugs

Ethanol 99.9% (containing less than 0.0002% benzene) was obtained from Merck. Src inhibitor 1 (SKI), sulfasalazine, tamoxifen, diphtheria toxin and rapamycin from Streptomyces hygroscopicus were from Sigma. AZD0530, pomalidomide and adalimumab were from Selleckchem.

### Alcohol intake protocol

Mice (genotypes are specified in the results and in the figure legends) were habituated for 4 weeks in experimental rooms at the i3S animal facility. Afterwards, mice were randomly assigned to experimental groups. To emulate a binge-like pattern of moderate alcohol intake, 1.5 g/Kg ethanol (diluted to 25% in sterile tissue culture-grade water) was administered by oral gavage daily for ten consecutive days. Mice were provided with food and water ad libitum throughout the experiments. Vehicle control mice were dosed with water in equal amounts to those of ethanol-treated mice. For analyses, mice were sacrificed 120 min after the last gavage. Measurements of blood alcohol concentration in adult C57BL/6 mice show that 120 min after oral intake of 1.5 g/Kg, ethanol is detected above 80 mg/dl in their blood [46, 47], thereby indicating a binge-like pattern of ethanol intake. For the in vitro studies, microglial cultures were exposed to 70 mM (320 mg/dl) ethanol. This concentration of ethanol was used because in addition to being correlated with a dosage inducing neuronal damage [53] it also simulates the amounts achieved in the blood during binge drinking.

### Pharmacological treatments

Cx3cr1^EYFP-CreER/+^ and WT mice were subjected to a simultaneous regimen of AZD0530 (Src inhibitor; 10 mg/kg) or DMSO (control) injections (pre-treatment intraperitoneal (IP) injection followed by 5 IP injections spaced by 24 hours during the ten-day ethanol exposure regimen). The same regimen was used for the immunomodulatory agent pomalidomide (PMD; 50 mg/kg). Tamoxifen was given to adult Cx3cr1^EYFP-CreER/+^ and Cx3cr1^EYFP-CreER/+^:R26^iDTR/+^ mice as a solution in corn oil by oral gavage. Mice received two doses of 10 mg of tamoxifen separated by 48 h between doses. For microglia ablation, 8 weeks after the last tamoxifen pulse, Cx3cr1^EYFP-CreER/+^ and Cx3cr1^EYFP-CreER/+^:R26^iDTR/+^ mice were given diphtheria toxin (1 µg; IP) for 3 consecutive days.

### Brain tissue preparation and immunohistochemistry

After animal perfusion with ice-cold PBS (15 ml) and fixation by perfusion with 4% PFA, brains were post-fixed by immersion in 4% PFA in PBS, pH 7.2 overnight. After that, brains were washed with PBS and then cryoprotected using sucrose gradient in a row (15 and 30%). After 24 h, brains were mounted in OCT embedding medium, frozen and cryosectioned in the CM3050S Cryostat (Leica Biosystems). Coronal sections from brains (30 μm thickness) were collected non-sequentially on Superfrost ultra plus slides. Tissue sections from controls and experimental mice encompassing identical stereological regions were collected on the same glass slide and stored at −20°C. Frozen sections were defrost by at least 1 h and hydrated with PBS for 15 min. Sections were permeabilized with 0.25% Triton X-100 for 15 min, washed with PBS for 10 min and blocked (5% BSA, 5% FBS, 0.1% Triton X-100) for 1 h. Primary antibodies were incubated in blocking solution in a humidified chamber overnight at 4°C. Secondary antibodies were incubated for 2 h in blocking solution. After the secondary antibody, sections were washed three times for 10 min with PBS. Slides were cover slipped using glycergel or Immumount and visualized under a Leica TCS SP5 II confocal microscope.

### Confocal imaging and morphometric analysis

Images from tissue sections of the prefrontal cortex and CA1 region of the dorsal hippocampus were acquired using a Leica HC PL APO Lbl. Blue 20x /0.70 IMM/CORR or a Leica HC PL APO CS 40x/1.10 CORR water objective in 8-bit sequential mode using standard TCS mode at 400 Hz and the pinhole was kept at 1 airy in the Leica TCS SP5 II confocal microscope. Images were resolved at 1024 x 1024 pixels format illuminated with 2-5% DPSS561 561 nm wave laser using a HyD detector in the BrightR mode and entire Z-series were acquired from tissue sections. Equivalent stereological regions were acquired for all tissue sections within a given slide.

#### Microglia quantification

Number of YFP^+^ cells was manually scored in stereological identical regions of the prefrontal cortex of stained sections (6 images per section; 5 sections per animal for each experimental group).

#### GFAP quantification

Number of GFAP^+^ cells was manually scored in stereological identical regions of the prefrontal cortex of stained sections (6 images per section; 8 sections per animal for each experimental group).

#### NeuN quantification

Number of NeuN^+^ neurons was manually scored in stereological identical regions of the prefrontal cortex stained sections (6 images per section; 8 sections per animal for each experimental group).

#### TNF quantification

Briefly, stereological identical regions of TNF immunostained sections of the prefrontal cortex or of the dorsal hippocampal CA1 region (4 images per section; 6 sections per animal for each experimental group) were imaged, converted into 8-bit gray scale, 3D volume-rendered and thresholded. Using FIJI software, the percent of TNF immunostained area was calculated for each field and each section.

#### Quantification of synapses

Images from stereological identical prefrontal cortex or dorsal hippocampal CA1 region from each experimental group (4 images per section; 6 sections per animal for each experimental group) were acquired using a Leica HC PL APO CS 40x /1.10 CORR water objective at 1024 x 1024 pixels resolution with 8-bit bidirectional non-sequential scanner mode at 400 Hz and pinhole at 1 airy in the Leica TCS SP5 II confocal microscope. Z-stacks were converted to maximum projection images using LAS AF routine. Using FIJI software, Z-projections were background subtracted and interpolated using a bicubic filter routine. The number of double positive PSD-95/vGlut1 puncta per µm^2^ was manually scored for each image.

#### Quantification of PSD-95 engulfment by microglia

Images from stereological identical prefrontal cortex or dorsal hippocampal CA1 region from each experimental group (6 images per section; 6 sections per animal for each experimental group) were acquired using a Leica HC PL APO CS 40x /1.10 CORR water objective at 1024 x 1024 pixels resolution with 8-bit bidirectional scanner mode at 200 Hz in the Leica TCS SP5 II confocal microscope. Using FIJI software, confocal Z stacks were background subtracted and smoothened using a Sigma filter plus. PSD-95 and Iba1 volumes were reconstructed using 3D surface rendering of confocal Z stacks in Imaris. For quantification of PSD-95 engulfment by microglia, PSD-95 puncta partially or completely embedded in volume-rendered Iba1^+^ structures were considered to be engulfed by microglia. The number of engulfed PSD-95 puncta was manually scored for each image and used for the analyses.

#### Assessment of microglia morphology

Ramification of prefrontal cortex Iba1+ microglia was assessed as described before [54]. Branches were traced using NeuronJ plugin, and data for each individual cells were converted into SWC format using Bonfire. Individual segments were connected in NeuronStudio software, and audited for any tracing errors using Bonfire. Sholl analysis was performed by drawing concentric circles around the cell body at defined radius increments, separated by 6 μm. The number of intersections of branches at each defined circle was used to estimate the number of ramifications. Branching data were extracted using a custom MATLAB routine.

### Flow cytometry

For characterization of immune cells in the samples, the following markers were used: CD45-PE (BioLegend 103106), CD11b-Alexa647 (BioLegend 101218), CD11b-APC (BioLegend 101212), CD68-BV421 (BioLegend 137017), CD3-APC-Cy7 (BioLegend 100221) and CD19-APC (BioLegend 115511). In brief, mice were anesthetized and then perfused with ice-cold PBS. For single cell suspensions, neocortices were quickly dissected on ice, placed on ice-cold RPMI and mechanically homogenized. Cells were collected, pelleted, washed extensively and then counted in a Neubauer chamber using trypan blue exclusion to estimate the number of live cells. Single cell suspensions (1 x 10^6^ cells) were incubated with different mixes of FACS antibodies for 30 min at 4°C in the dark. Compensation settings were determined using spleens from wild-type mice. Cell suspensions were evaluated on a FACS Canto II analyzer (BD Immunocytometry Systems). Microglial cell numbers found on neocortices were estimated using Precision Count Beads^TM^ (BioLegend 424902).

### Preparation of lysates and Western blotting

Cultures or mice tissues were lysed using RIPA-DTT buffer (150 mM NaCl, 50 mM Tris, 5 mM EGTA, 1% Triton X-100, 0.5% DOC, 0.1% SDS) supplemented with complete-mini protease inhibitor cocktail tablets, 1 mM DTT and phosphatase inhibitor cocktail. Samples were sonicated (6 pulses of 1 sec at 60Hz) and centrifuged at 16,000 g, 4°C for 10 min. The supernatants were collected and the protein concentration was determined by the BCA method. All samples were denatured with sample buffer (0.5 M Tris-HCl pH 6.8, 30% glycerol, 10% SDS, 0.6 M DTT, 0.02% bromophenol blue) at 95°C for 5 min and stored at −20°C until use. Samples were separated in SDS-PAGE, transferred to PVDF membranes, which were incubated overnight with primary antibodies. Membranes were washed in TBS-T buffer pH 7.6, incubated with peroxidase-conjugated secondary antibodies and developed using an ECL chemiluminescence kit or an ECF fluorescence kit. Images were acquired in a Typhoon FLA 9000 system or ChemiDoc XRS System (Bio-Rad) and quantified by FIJI software.

### Gene expression and bioinformatic analyses

RNA was extracted from neocortices using the Direct-zol^TM^ RNA MiniPrep kit according to the manufacturer’s instructions. cDNA synthesis was performed using 500 ng of total RNA (DNase I treated) with SuperScript^®^ III First-Strand Synthesis SuperMix. qRT-PCR was carried out using iQ™ SYBR^®^ Green Supermix on an iQ™5 multicolor real-time PCR detection system (Bio-Rad). Expression of PCR transcripts was calculated using the 2^-deltaCt^ with *Yhwaz* serving as the internal control gene. Statistical analysis on raw 2^-deltaCt^ values using unpaired t tests were performed for detecting differentially expressed transcripts between sampled groups. For representation, transcript expression values were converted into Z scores. ClueGo [55] was used for unsupervised enrichment analyses and statistical overrepresentation of immune signaling pathways. The NetworkAnalyst 3.0 [56] was used for constructing topographic maps of overrepresented pathways from gene ontology (PANTHER/SLIM GO) and for generating PPI networks (using the STRING database for representation of interacting protein clusters).

### Behavioral tests

All testing procedures were conducted in the dark phase of the light/dark cycle. Before each session, mice were removed from their home cage in the colony room and brought into the adjacent testing rooms (illuminated with 100 lux and attenuated noise). Behavioral tests were performed in the following order: (1) elevated plus-maze; (2) open field; (3) novel object recognition.

#### Elevated plus-maze (EPM)

The maze was made of opaque grey PVC consisting of four arms arranged in a plus-shaped format; two arms have surrounding walls (closed arms, 37×6 cm x18 cm-high), while the two opposing arms have no walls (open arms, 37×6 cm). The apparatus is elevated by 50 cm above the ground. Mice were placed on the central platform facing an open arm and were allowed to explore the maze for 5 minutes. Frequency and time spent in the open arms were obtained automatically (Smart 3.0 Panlab Harvard Apparatus) and used to assess anxiety-like behavior.

#### Open field (OF)

Mice were placed in the center of an OF apparatus (40 x 40 x 40 cm) and then allowed to move freely for 10 min. The total distance travelled, as well as locomotion in the peripheral zone and center zone of the apparatus were obtained automatically using video tracking.

#### Novel object recognition (NOR)

The NOR test was performed as previously described [57]. Briefly, the same OF test apparatus was employed and the objects used were made of plastic, glass or metal in three different shapes: cubes, pyramids and cylinders. The test consists of three phases. During habituation phase mice are allowed to explore the apparatus for 10 min (period considered for the OF test). The following day, the acquisition/sample phase starts by placing each mouse in the apparatus with two identical objects (familiar) for 10 min. Then the mouse goes back to its home cage. After 4 h (inter-trial interval, ITI), the retention/choice session is performed. In this phase, the apparatus contains a novel object and a copy of the previous familiar object; animals are allowed to explore these objects for 3 min. Exploration was defined as follows: mouse touched the object with its nose or the mouse’s nose was directed toward the object at a distance shorter than 2 cm [58]. Circling or sitting on the object was not considered exploratory behavior. Behavioral data were collected using the software Observer XT 7.0, Noldus. Increased time spent exploring the novel object serves as a measure of recognition memory for the familiar object. The discrimination index (DI) was calculated as index of memory function, DI = (time exploring the novel object - time exploring the familiar object) / (total time spent exploring both objects). Positive values indicate increased time investigating the novel object, which serves as indication of object discrimination.

### Primary cortical microglial cultures

Primary cortical microglial cell cultures were performed as previously described [19–21]. In brief, mice pups (2-day-old) were sacrificed, their cerebral cortices were dissected in HBSS, pH 7.2, and digested with 0.07% trypsin plus 50 μL (w/v) DNAse for 15 min. Next, cells were gently dissociated using a glass pipette in DMEM F12 GlutaMAX™-I supplemented with 10% FBS, 0.1% gentamicin. Cells were plated in poly-D-lysine-coated T-flasks (75 cm^2^) at 1.5×10^6^ cells per cm^2^. Cultures were kept at 37°C and 95% air/5% CO_2_ in a humidified incubator. Culture media was changed every 3 days up to 21 days. To obtain purified microglial cell cultures, culture flasks were subjected to orbital shaking at 200 rpm for 2 h. Next, culture supernatant was collected to tubes, centrifuged at 453 g for 5 min at room temperature. The supernatant was discarded and the pellet, containing microglia, was re-suspended in culture medium, and cells were seeded in poly-D-lysine-coated 6 or 12-well culture plates at 2.5 x 10^5^ cells/cm^2^ with DMEM F12 GlutaMAX™-I supplemented with 10% FBS, 0.1% gentamicin and 1 ng/ml GM-CSF. Purified microglia were cultured for 5-8 days. Immunolabeling with CD11b showed a purity of 95-99% for these cultures.

### Microglial cell lines

The microglial cell line N9 was obtained by immortalization of primary cultures from the ventral mesencephalon and cerebral cortex from ED12-13 CD1 mouse embryos with the 3RV retrovirus carrying an activated v-myc oncogene [59]. The microglial cell line CHME3 was obtained from primary cultures of human embryonic microglial cells by transfection with a plasmid encoding for the large T antigen of SV40 [60]. Cells were cultivated and maintained as before [19].

### Synaptosomal preparations and microglia engulfment assay

To isolate prefrontal cortex synaptic terminals synaptosomes were freshly prepared using Syn-PER™ Synaptic Protein Extraction Reagent (catalog no. 87793, ThermoFisher Scientific) exactly as recommended by the manufacturer. Briefly, WT mice or Thy1-YFP were euthanized in a CO_2_ chamber. Their prefrontal cortex was collected and homogenization was performed using a Dounce tissue grinder (∼10 strokes). The homogenate was centrifuged at 1200 x *g* for 10 min and the pellet was discarded. The supernatant was centrifuged at 15000 x *g* for 20 min. The resulting supernatant was discarded and the synaptosomal pellet was resuspended in Syn-Per™ Synaptic Protein Extraction Reagent.

CHME3 microglial cultures were seeded in 12-well culture plates at a density of 2.5 x 10^4^ cells/well in DMEM GlutaMAX™-I supplemented with 10% FBS, 0.1% PenStrep and cultivated for 48 h. Cultures were then refed with fresh medium containing vehicle (water), or EtOH (70 mM), or adalimumab (5 μg/ml), or adalimumab (5 μg/ml) plus EtOH (70 mM). Cells were kept in those conditions for 24 h. Afterwards, cells were refed with fresh medium containing vehicle (water) plus synaptosomes (1:100), or EtOH (70 mM) plus synaptosomes (1:100), or adalimumab (5 μg/ml) plus synaptosomes (1:100), or adalimumab (5 μg/ml) plus EtOH (70 mM) plus synaptosomes (1:100). Cells were cultured under these conditions for additional 24 h. For analyses, microglial cultures were washed extensively with PBS and either fixed with PFA 4%, immunolabeled for PSD-95 and analyzed in a fluorescence microscope or detached using accutase solution (catalog no. A6964, Sigma), blocked in PBS-BSA 5%, fixed with PFA 2%, resuspended in PBS-BSA 1% and analyzed in a flow cytometer.

### Phagocytic assay in primary cortical microglia

Cell cultures were incubated with fluorescent latex microbeads (0.001% v/v final concentration; Sigma) for 90 min. Cultures were washed 3x with PBS and fixed in 4% PFA (w/v) for 12 min. Afterwards, f-actin was stained using phalloidin and cells were visualized in a fluorescence microscope. The number of engulfed microbeads per microglia was manually scored and plotted for statistical evaluation.

### Plasmids

NFκB GFP–tagged p65 (Plasmid 23255), IκBα-miRFP703 (Plasmid 80005), pLNCX chick src Y527F (plasmid 13660), pUSE-RapR-Src-myc (plasmid 25933), psPAX2 (plasmid 12260), pMD2.G (plasmid 12259), pUMVC (plasmid 8449), pMSCV (plasmid 24828), and pCherry-FRB (plasmid 25920) were from Addgene.

### Retroviruses production

Low passage HEK293T cells were seeded in 100 mm culture dishes. When cultures reached ∼80% confluence cells were co-transfected overnight with viruses-producing plasmids using the transfection reagent jetPRIME^®^. Transfection ratios were as follows: 6 μg of shRNA plasmids to 3 μg of psPAX2 to 3 μg of VSVG (2:1:1) for lentiviruses production or 8 μg of Src^Y527F^ construct to 4 μg of pUMVC to 2 μg of VSVG (4:2:1) for viral production. The next day, normal growth media replaced transfection media and cells were cultivated for an additional 48 h. Next, media with viral particles were collected, centrifuged at 906 g for 15 min at 4°C, and the supernatant was collected into new tubes and kept at −80°C.

### Live cell imaging and FRET

Microglial cells were plated on plastic-bottom culture dishes (µ-Dish 35 mm, iBidi). Imaging was performed using a Leica DMI6000B inverted microscope. The excitation light source was a mercury metal halide bulb integrated with an EL6000 light attenuator. High-speed low vibration external filter wheels (equipped with CFP/YFP/farRed excitation and emission filters) were mounted on the microscope (Fast Filter Wheels, Leica Microsystems). A 440-520nm dichroic mirror (CG1, Leica Microsystems) and a PlanApo 63X 1.3NA glycerol immersion objective were used for CFP and FRET images. Images were acquired with 2×2 binning using a digital CMOS camera (ORCA-Flash4.0 V2, Hamamatsu Photonics). At each time-point, CFP, FRET and fared images were sequentially acquired using different filter combination. [19–21]. For quantifications, images were exported as 16-bit tiff files and processed in FIJI software. Background was dynamically subtracted from all frames from both channels. Segmentation (on a pixel-by-pixel basis) and generation of 32-bit float-point ratiometric images was achieved using the precision FRET (PFRET) data processing software package for ImageJ (https://lvg.virginia.edu/digital-downloads/pfret-data-processing-software). The mean gray intensity values from ratio images were used for statistical calculations.

### Cytokine release

Culture medium was collected to tubes and centrifuged at 16,000 g at 4°C for 5 min. The supernatant was transferred to a new tube and kept at −80°C. The concentration of TNF in cell culture supernatants was quantified by enzyme-linked immunosorbent assay (ELISA) following the instructions provided by the manufacturer (Peprotech, UK). Absorbance at 405 nm, with wavelength correction at 650 nm, was measured with a multimode microplate reader (Synergy HT, Biotek, USA). Values corresponding to ng/ml were obtaining by extrapolating a standard concentration curve using recombinant TNF.

### Statistics

A 95% confidence interval was used for statistical evaluation and p < 0.05 was considered statistically significant difference in all sampled groups. Experimental units in individual replicates were prior evaluated for Gaussian distribution using the D’Agostino & Pearson omnibus normality test with GraphPad Prism. When comparing only 2 experimental groups Mann-Whitney test for data with non-normal distribution or unpaired Student t test with equal variance assumption for data with normal distribution was used. When comparing 3 groups, One-way ANOVA followed by the Fisheŕs multiple comparation test was used. When comparing 4 or more groups, Two-way ANOVA followed by the Turkeýs multiple comparation test was used. Two-way ANOVA was also used to compare values of microglia intersections retrieved from Sholl analysis. To minimize bias, experimental groups were assigned through randomization. Specifically, sampled groups evaluated in Figures 1, 2 and in all Supplemental Figures were assigned using ‘simple random sampling’. Sampled groups evaluated in Figures 3, 4, 5 and 6 were assigned using ‘permuted block randomizatio’. All quantifications were performed blinded.

## Supporting information

Primers list

