## Supplementary material for "Alcohol intake triggers aberrant synaptic pruning leading to synapse loss and anxiety-like behavior": Suppl Legends

**Supplemental Legends**

**S1.** Alcohol exposure does not activate other immune resident cells in the neocortex. (Related to Fig. 1).

**A**, flow cytometry analysis of microglial numbers in neocortices of Cx3cr1^EYFP-CreER/+^ and WT mice exposed to EtOH or H_2_O (n=5 animals per genotype). Graphs (mean and SEM) show microglia cell counts. Cell debris was excluded by size. *P<0.05 (One-Way ANOVA vs. Cx3cr1^EYFP-CreER/+^ H_2_O); ^†^p<0.05 (One-Way ANOVA vs. WT H_2_O).

**B,** qRT-PCR from neocortices of Cx3cr1^EYFP-CreER/+^ and WT mice exposed to EtOH or H_2_O (n=5 animals per genotype). Graph (mean and SEM) shows the mRNA expression for the indicated transcripts. *P<0.05 (One-Way ANOVA vs. Cx3cr1^EYFP-CreER/+^ H_2_O); ^†^p<0.05 (One-Way ANOVA vs. WT H_2_O).

**C**, flow cytometry analysis of macrophages, B lymphocytes and T lymphocytes in neocortices of Cx3cr1^EYFP-CreER/+^ mice exposed to EtOH or H_2_O (n=8 animals per group). Graphs (mean and SEM) show cell counts. Cell debris was excluded by size. Statistical comparations were performed using Mann-Whitney test.

**D,** histological confocal analysis of astrocyte number on tissue sections from prefrontal cortices of Cx3cr1^EYFP-CreER/+^ mice exposed to EtOH or H_2_O (n= 5 animals per group). Graph (mean and SEM) shows the number of GFAP+ cells. Statistical comparations were performed using Mann-Whitney test. Scale bar: 50 μm.

**E**, Western blot for GFAP on lysates from prefrontal cortices of Cx3cr1^EYFP-CreER/+^ mice exposed to EtOH or H_2_O (n=5 animals per group). Graph (mean and SEM) shows normalized ratio of GFAP over GAPDH (loading control). Statistical comparations were performed using Mann-Whitney test.

**S2.** Alcohol exposure modulates microglial TNF production via Src/NFkB pathway. (Related to Fig. 4).

**A,** primary cortical microglial cultures were treated with EtOH (70 mM) or H_2_O (n=14 different and independent cultures) for 24 h and immunostained for phospho-Src Tyr^416^ (red). Graph (mean and SEM) displays the normalized pSrc fluorescence intensity per individual microglia in each cell culture. *P<0.05 (Mann-Whitney test).

**B**, CHME3 microglial cells expressing a Src FRET biosensor (KRas Src YPet) were recorded under basal condition (in the presence of HBSS) and then were challenged with 70 mM EtOH (n=25 cells pooled across 5 independent experiments). CFP/FRET emission ratios of the biosensor were normalized at 0 min. The panels show time-lapse CFP/FRET images coded according to the indicated pseudocolor scale. Graph displays the normalized maximum CFP/FRET amplitude in EtOH vs. HBSS. *P<0.001 (Mann-Whitney test). Scale bar: 10 μm.

**C**, Western blot for phospho-Src Tyr^416^ and Src on lysates from neocortices of Cx3cr1^EYFP-CreER/+^ mice treated with DMSO or AZD0530 (gels are representative of 5 different animals per group).

**D,** primary cortical microglial cultures were treated with EtOH (70 mM) or H_2_O (n=6 different and independent cultures) for 24 h and immunostained for TNF. Graph (mean and SEM) displays the normalized TNF fluorescence intensity per individual microglia in each cell culture coded according to the indicated pseudocolor scale. *P<0.05, ^§^not significantly different (One-way ANOVA). Scale bar: 10 μm.

**E**, CHME3 microglial cells expressing a GFP-tagged p65 NFkB subunit (p65-GFP) were treated for 24 h with EtOH (70 mM) or H_2_O (n=8 different and independent cultures). Graph displays the nuclear GFP fluorescence intensity per cell. *P<0.05, ^§^not significantly different (One-way ANOVA). Scale bar: 10 μm.

**F**, Western blot for phospho-Src Tyr^416^ and Src on lysates from CHME3 microglial cells expressing a vector coding for Src shRNA or a control (pLKO) vector (gels are representative of 4 different cultures).

**G**, CHME3 microglial cells expressing a NFkB pathway inhibitor biosensor were recorded under basal condition (in the presence of HBSS) and then were challenged with 70 mM EtOH (n=15 cells per group pooled across 3 independent experiments). Fluorescence signal of time lapses was normalized at 0 min in each experimental group. *P<0.05 vs. EtOH 0 min; ^§^P<0.05 AZD + EtOH vs. EtOH at each related time point; ^†^P<0.05 SKI-1 + EtOH vs. EtOH at each related time point (Two-way ANOVA).

**H**, CHME3 microglial cells co-expressing a GFP-tagged p65 NFkB subunit (p65-GFP) with Src chemogenetic constructs (FRB + RapR-Src) or control constructs (FRB + FKBP) were treated with rapamycin (200 nM) and EtOH (70 mM) or H_2_O for 24 h (n=12 different and independent cultures). Graph displays the nuclear GFP fluorescence intensity per cell. *P<0.05 (Mann-Whitney test). Scale bar: 20 μm.

**I**, ELISA on the culture media of N9 microglial cultures treated for 24 h with EtOH (70 mM) or H_2_O (n=4 different and independent cultures). Graph (means and SEM) displays TNF content in ng/ml. *P<0.05 (Mann-Whitney test).

**J,** N9 microglial cultures expressing an empty pMSCV vector (EV) or the Src^Y527F^ mutant were treated with 1 mM sulfasalazine for 24 h (n=4-6 different and independent cultures) and immunostained for TNF and CD11b. Graph (mean and SEM) displays the normalized TNF fluorescence intensity per individual microglia in each cell culture coded according to the indicated pseudocolor scale. *P<0.05, ^§^ P<0.05 (One-way ANOVA). Scale bar: 20 μm.

**S3.** Alcohol exposure does not alter general locomotor activity, recognition memory or prefrontal cortex neuronal cell numbers. (Related to Fig. 5).

**A and B**, WT mice were exposed to EtOH or H_2_O and then evaluated in the elevated-plus maze (EPM) test (n=7 animals per group). Graphs (mean and SEM) show the frequency of entries in the open arms (A) and total distance animals travelled in the maze (B). *P<0.05 (Mann-Whitney test).

**C-E**, Cx3cr1^CreER/+^ mice were exposed to EtOH or H_2_O and then evaluated in the open field arena (n=8 animals per group). Graphs are mean and SEM and statistical comparations were analyzed by Mann-Whitney test.

**F and G**, Cx3cr1^CreER/+^ mice were exposed to EtOH or H_2_O and then evaluated in the novel object recognition (NOR) test (n=8 animals per group). Recognition memory was similar between genotypes. Graphs are mean and SEM and statistical comparations were analyzed by Mann-Whitney test.

**H,** histological confocal analysis of NeuN on tissue sections from prefrontal cortices of Cx3cr1^EYFP-CreER/+^ mice exposed to EtOH or H_2_O (n=5 animals per group). Graph (mean and SEM) shows the number of NeuN+ cells. Statistical comparation was performed using Mann-Whitney test. Scale bar: 50 μm.

**S4.** Alcohol exposure does not alter synapse number, microglia postsynaptic engulfment or TNF production in the CA1 region of the dorsal hippocampus. (Related to Fig. 5).

**A,** histological confocal colocalization analysis for PSD-95 (red) and vGlut1 (green) on tissue sections from the CA1 region of the dorsal hippocampus of Cx3cr1^EYFP-CreER/+^ mice after exposure to EtOH or H_2_O (n=6 animals per group). Graph (mean and SEM) shows the normalized number of excitatory synapses (yellow) and statistical comparation was analyzed by Mann-Whitney test. Scale bar: 5 μm.

**B,** histological confocal colocalization analysis for PSD-95 (red) and Iba1 (green) on tissue sections from the CA1 region of the dorsal hippocampus of Cx3cr1^EYFP-CreER/+^ mice after exposure to EtOH or H_2_O (n=6 animals per group). Graph (mean and SEM) shows the normalized number of PSD-95 puncta engulfed by microglia (yellow) and statistical comparation was analyzed by Mann-Whitney test. Scale bar: 10 μm.

**C,** histological confocal analysis for TNF on tissue sections from the CA1 region of the dorsal hippocampus of Cx3cr1^EYFP-CreER/+^ after exposure to EtOH or H_2_O (n=5 animals per group). Graph (mean and SEM) shows the TNF protein expression coded according to the pseudocolor ramp. Statistical comparation was analyzed by Mann-Whitney test. Scale bar: 100 μm.
