## Supplementary figures and images for "Alcohol intake triggers aberrant synaptic pruning leading to synapse loss and anxiety-like behavior"

### Suppl. Figure 1

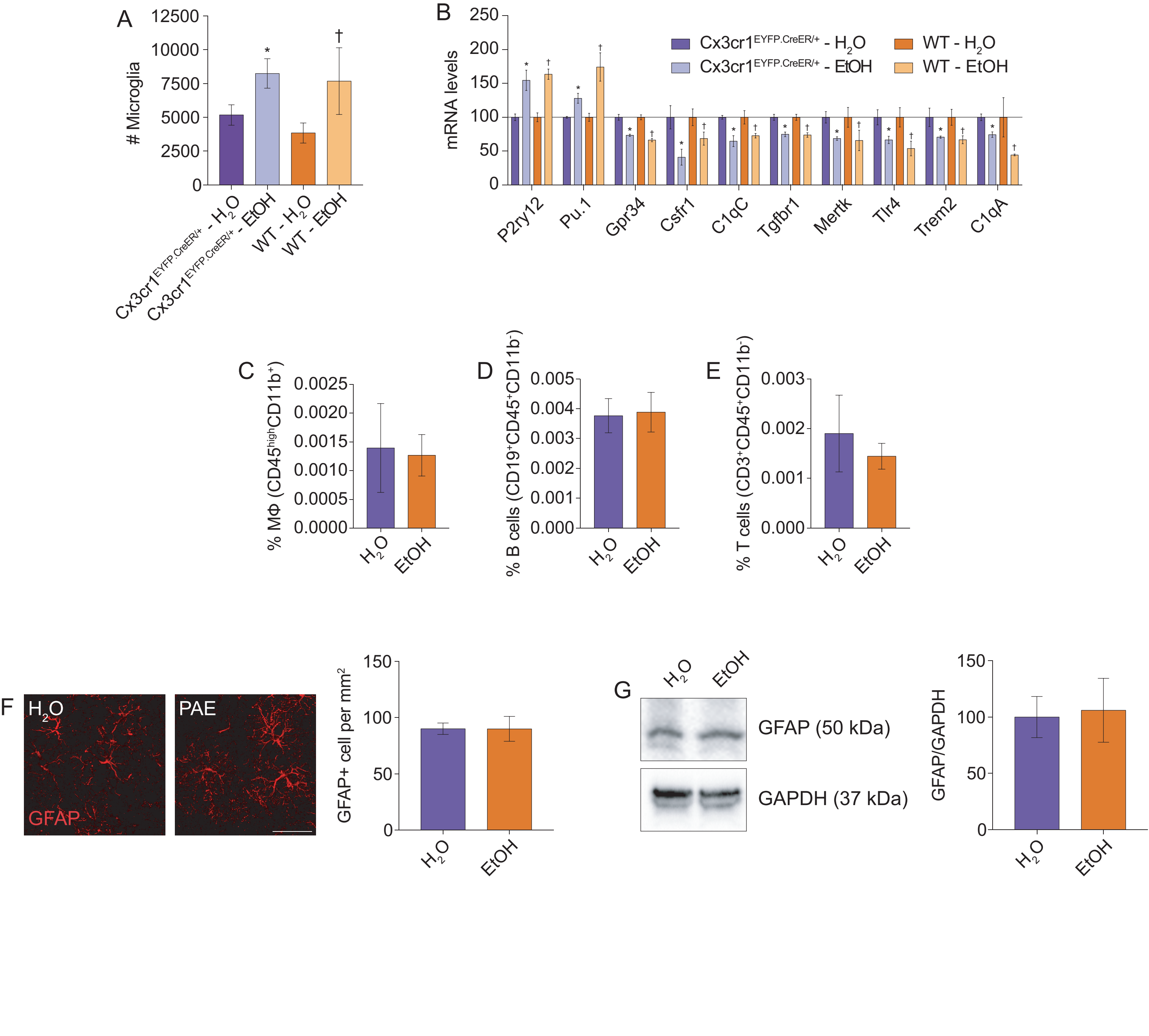

### Suppl. Figure 2

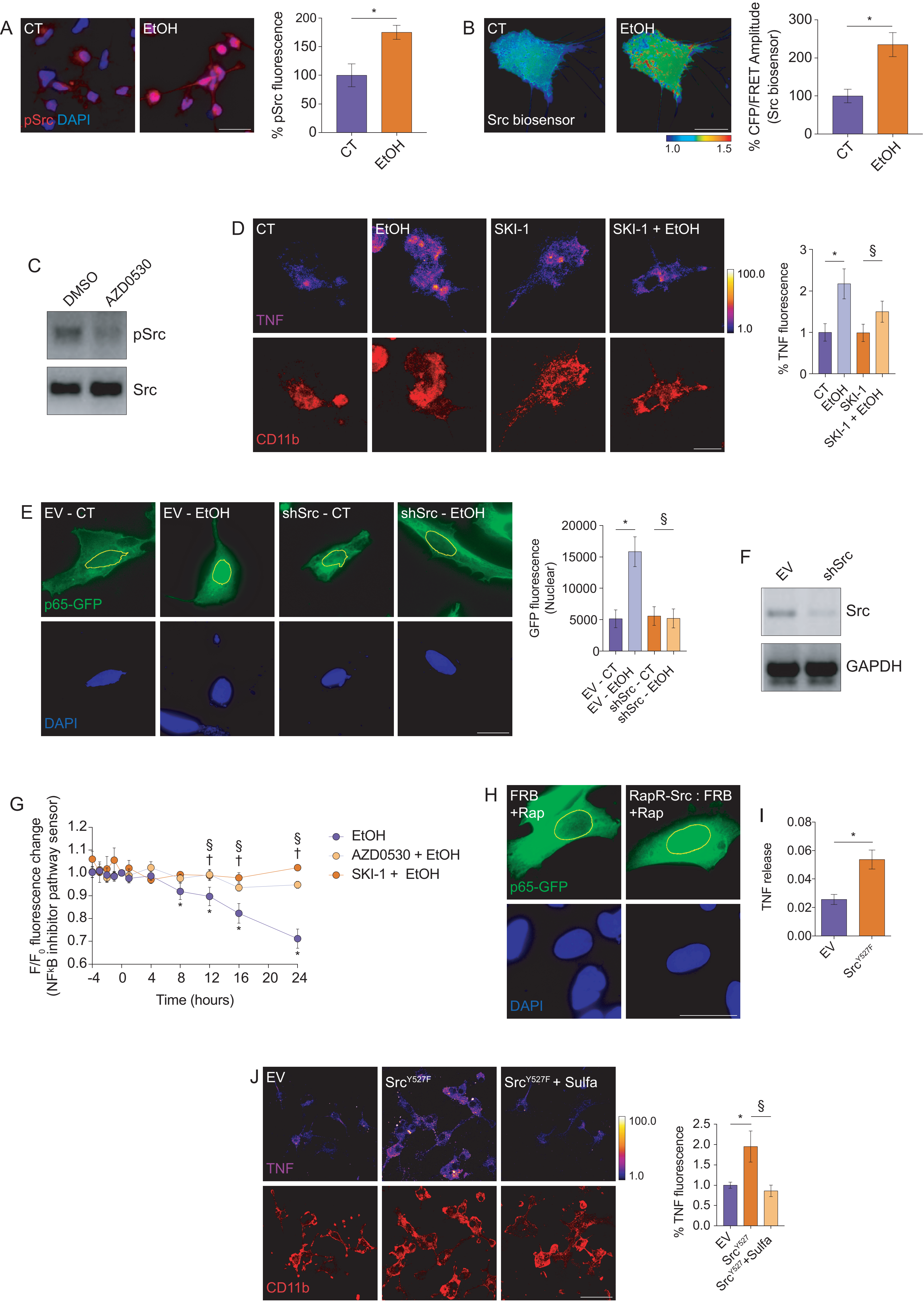

### Suppl. Figure 3

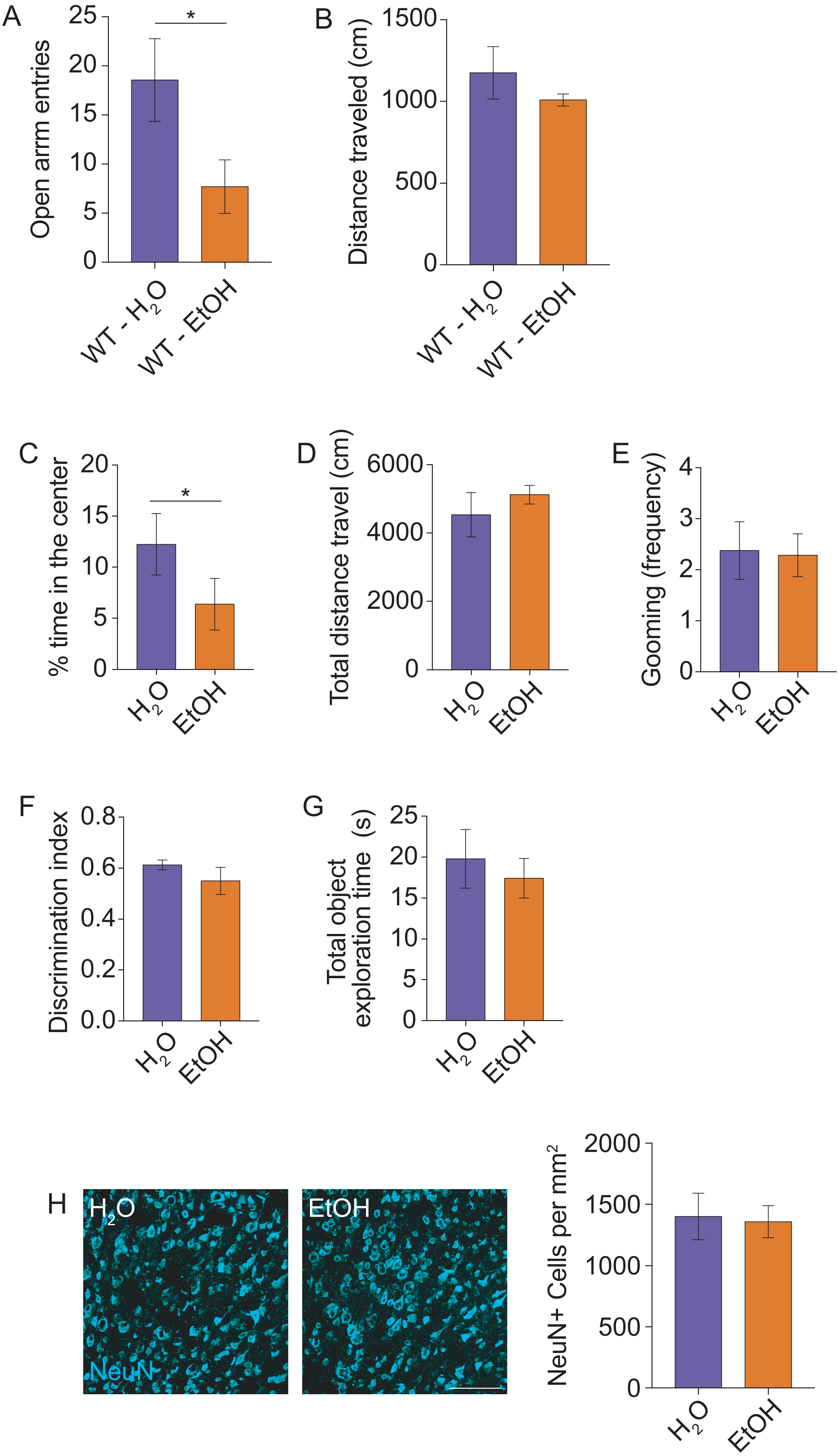

### Suppl. Figure 4

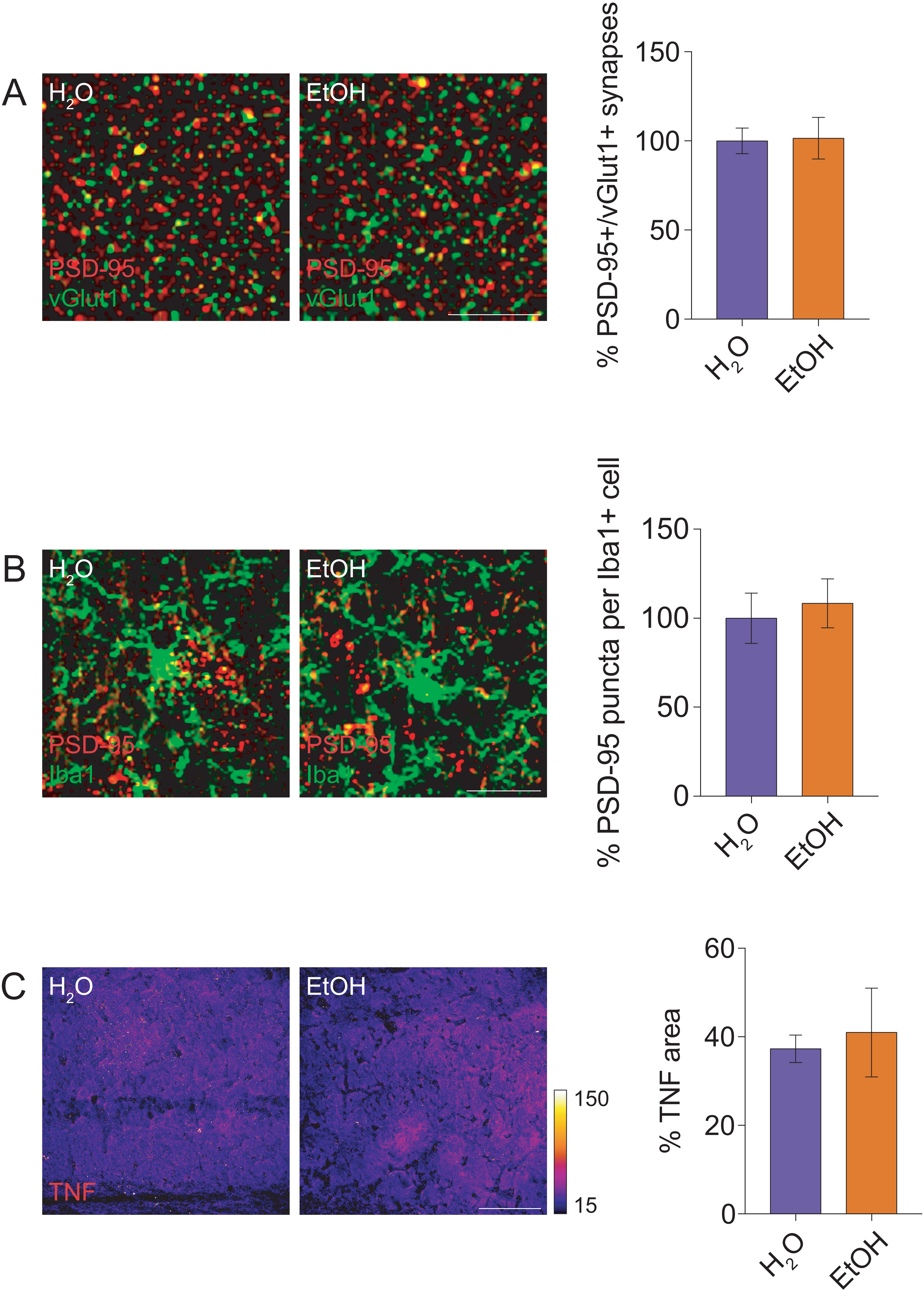
